# The Adr1 transcription factor directs regulation of the ergosterol pathway and azole resistance in C. *albicans*

**DOI:** 10.1101/2023.01.24.524502

**Authors:** Manjari Shrivastava, Gaëlle S. Kouyoumdjian, Eftyhios Kirbizakis, Daniel Ruiz, Manon Henry, Antony T Vincent, Adnane Sellam, Malcolm Whiteway

## Abstract

Transcription factors play key roles in cellular regulation and are critical in the control of drug resistance in the fungal pathogen *Candida albicans*. We found that activation of the transcription factor C4_02500C_A (Adr1) conferred significant resistance against fluconazole. In *Saccharomyces cerevisiae*, Adr1 is a carbon-source-responsive zinc-finger transcription factor required for transcription of the glucose-repressed gene *ADH1* and of genes required for ethanol, glycerol, and fatty acid utilization. Motif scanning of promoter elements suggests that Adr1 may be rewired in fungi and governs the ergosterol synthesis pathway in *C. albicans*. Because previous studies have identified the zinc-cluster transcription factor Upc2 as a regulator of the ergosterol pathway in both fungi, we examined the relationship of Adr1 and Upc2 in sterol biosynthesis in *C. albicans*. Phenotypic profiles of either *ADR1* and *UPC2* modulation in *C. albicans* showed differential growth in the presence of fluconazole; either *adr1* or *upc2* homozygous deletion results in sensitivity to the drug while their activation generates a fluconazole resistant strain. The rewiring from ergosterol synthesis to fatty acid metabolism involved all members of the Adr1 regulon except the alcohol dehydrogenase Adh1, which remains under Adr1 control in both circuits and may have been driven by the lifestyle of *S. cerevisiae*, which requires the ability to both tolerate and process high concentrations of ethanol.

## Introduction

*Candida albicans* is an opportunistic fungal pathogen that is responsible for a variety of fungal infections in humans. In healthy people, this yeast resides as a commensal in the gastrointestinal tract, but it is capable of causing mucosal, cutaneous, and systemic infections in immunocompromised individuals [1]. The prevalence of resistance to antifungal agents has significantly increased in the past few decades, and this resistance has important implications for mortality, morbidity, and health care in the community [2]. The development of new antifungal drugs is challenging, as fungi are eukaryotic organisms that share many basic cellular processes with humans, and this evolutionary relatedness makes the identification of specific targets difficult and increases the likelihood of undesired secondary effects. Existing antifungals tend to target processes that are divergent between fungi and the human host.

The azole class of antifungals, including fluconazole, targets the ergosterol pathway, inhibiting a step not found in the pathway for the host-specific sterol cholesterol. Azoles are generally effective for the management of *C. albicans* infections, but due in part to the fungistatic nature of the drugs, long-term treatment often results in the emergence of azole resistance, ultimately resulting in therapeutic failure [3–5]. These azole antifungals bind and inhibit the activity of the enzyme lanosterol 14-alpha-demethylase encoded by *ERG11* [6]. Apart from azoles, allylamines (which target Erg1), polyenes (which target ergosterol itself), morpholines (which target Erg2) and statins (which target HMG1/2) also target elements of the sterol pathway [7]. As many drugs target the *C. albicans* sterol pathway, genetic changes that perturb the pathway can lead to multidrug resistance [8, 9].

A promising approach for drug development is to identify synergistic targets that can enhance the antifungal effect of currently available drugs [10]. Transcription factors (TFs) play a key role in determining how cells function and respond to different environments, and approximately 4% of *C. albicans* genes encode for transcription factors [11], making them the single largest family of proteins in the pathogen. TFs in *C. albicans* coordinate critical cellular functions including biofilm formation [12], drug resistance [13], and the transition from a commensal to a pathogenic lifestyle [14].

Most transcription factors are conserved, in that they fall into a limited number of groups of structurally similar proteins, such as the zinc finger, the basic helix loop helix, and the leucine zipper classes. However, evolutionary changes in transcription networks are an important source of diversity across species, driven primarily not by major changes in the structures of the factors themselves, but in the connections between the transcription factors and their regulated genes. There are many incidences where researchers have identified structurally equivalent transcription factors regulating different genetic circuits in different organisms [15–18]. This phenomenon has been called “rewiring” of transcription factors. Studies suggest that this rewiring happens at a relatively constant rate, and for two species that have diverged for 100 million years, only a fraction of transcription factor/target gene combinations will likely have remained conserved [19, 20]. *C. albicans* belongs to the same family as *Saccharomyces cerevisiae*, but the two fungi are suggested to have diverged as long ago as 300 million years, allowing for considerable rewiring. While studies of TFs have tended to focus on similarities between these two species, it has been estimated that only 16% of the regulator-target gene connections are preserved between the *C. albicans* and *S. cerevisiae* [21].

In our study we have activated a group of transcription factors of *C. albicans* for which there was limited information, and which had a potential of being rewired. Among our tested set of TFs, we found that C4_02500C_A activation gives resistance to several cell-membrane targeting drugs. This resistance arises because C4_02500C_A is a central regulator of the ergosterol pathway in *C. albicans*. Further analysis shows that this TF is the ortholog of *S. cerevisiae* Adr1, and that the two proteins play distinct cellular roles in the two species.

## Results

### Fusion of different transcription factors to the strong activation domain VP64

In *S. cerevisiae* the fusion of VP16 to the N terminus of Gal4 resulted in the hyper-activation of Gal4 (Daniel et al., 1998). VP64 fusion has been used to successfully activate transcription factors in both plants [23] and animals [24]. We have used a similar strategy in *C. albicans*. Fusing a tetrameric version of the VP16 trans-activating domain (VP64) to the DNA binding domains of different *C. albicans* transcription factors was found to be even more potent in transcriptional activation [22]. We constructed plasmid CIPACT-VPR containing the VP64 module and a multiple cloning site (MCS) downstream of the *ACT1* promoter of the CIPACT plasmid (Fig. S1A).

To test this strategy, we choose 3 well-studied transcriptional factors - the bZIP TF Gcn4 (null mutant gives 3 amino-triazole sensitivity), a TEA/ATTS (Homeo-domain) TF Tec1 (null mutant blocks hyphal development), and a leucine bZIP TF Cap1 (involved in fluconazole resistance). The Gcn4 construct generated resistance to 3 amino-triazole, consistent with up-regulation of the Gcn4 target *HIS3* (Fig. 1A). The Tec1 construct triggered filamentation under yeast morphology growth conditions (Fig. 1B), and the Cap1 activation construct enhanced resistance to fluconazole (Fig. 1C).

**Fig. 1.**
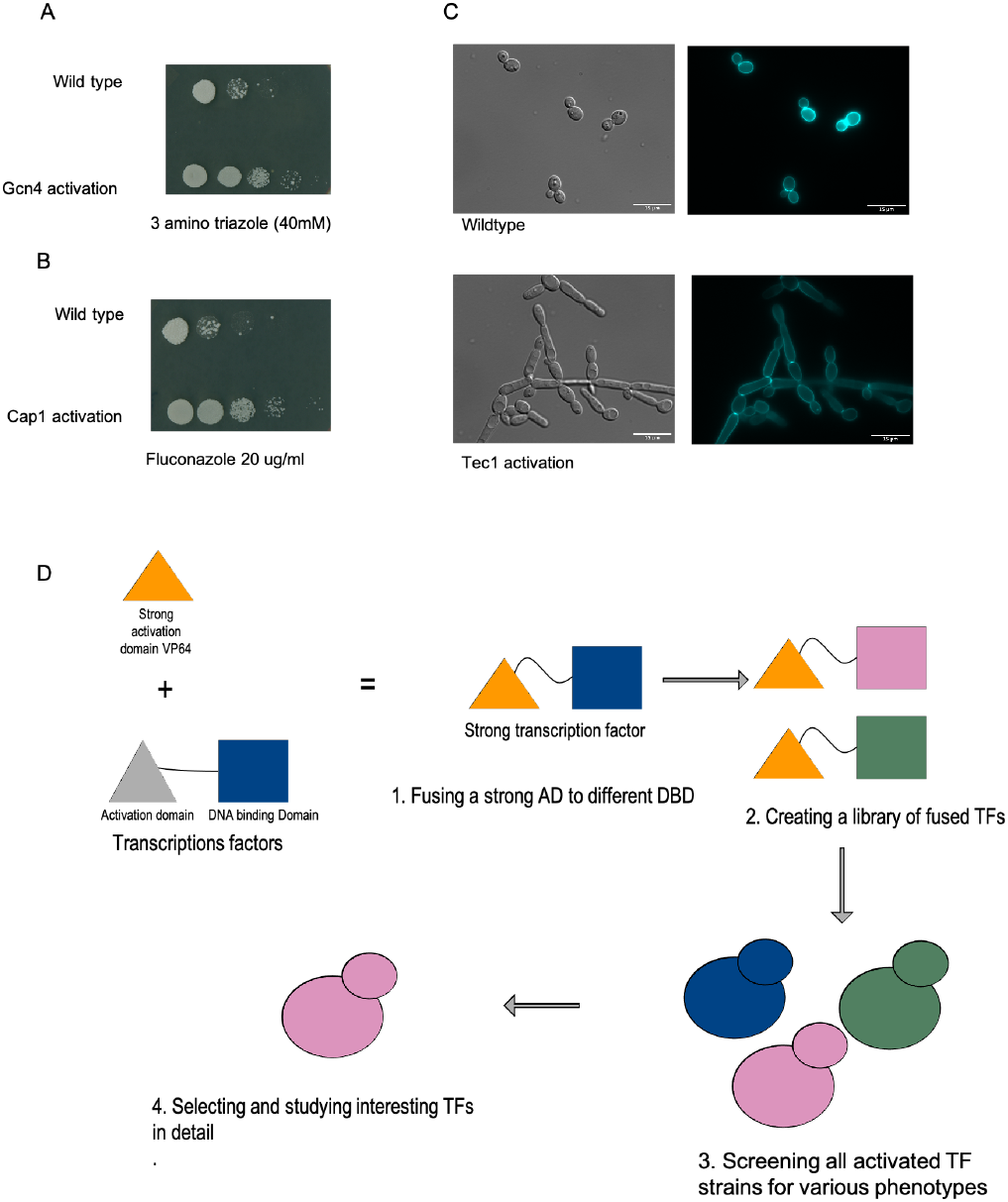
**(A). VP64-Gcn4 chimeric transcription factor generates resistance to 3-amino-triazole (3AT).** To construct the Gcn4-Vp64 fusion, we PCR amplified the Gcn4 DNA-binding domain and ligated it in the CIP-ACT-CYC plasmid in-frame with VP64 at the N terminus. After transforming this plasmid into C. albicans we observed resistance to 3AT consistent with the VP64 module activating the transcription factor. **(B) VP64-Tec1 TF triggers hyphal elongation in YPD media.** Tec1 is a transcription factor implicated in the morphological switch from the C. albicans yeast form to the hyphal form. A construct containing the fusion of the N terminus of Tec1 to the VP64 module triggers elongated cellular growth. **(C) VP64-Cap1 allows growth in SC media containing fluconazole.** Cap1 is a poorly characterized transcription factor in C. albicans that gives resistance to azoles through Mdr1; activation of Cap1 upregulates MDR1 expression. Fusion of the VP64 module to Cap1 increased cellular tolerance to the azole fluconazole. **(D) Schematic representation of the workflow involved in activating the transcription factors.** Based on the success of the control constructs, we selected a set of transcription factors for fusion constructs and characterized the consequences of these fusions through phenotypic analysis.

We next selected 30 different TFs based on their phylogenetic uniqueness, their possible involvement in drug resistance, and their potential of functioning in rewired circuits. After generating these 30 TF-VP64 fusions we first investigated their involvement in antifungal drug resistance (Fig. 1D). We selected 3 drugs for our preliminary screening - fluconazole, posoconazole, and amphotericin B. All three drugs target the cell membrane; fluconazole and posoconazole target lanosterol 14-alpha-demethylase (Erg11), an enzyme of the ergosterol pathway, whereas amphotericin B targets the end-product ergosterol. We identified two transcription factor fusions, encoded by C4_02500C_A and C6_00010W_A, which gave resistance to all three drugs. As the C4_02500C_A fusion created a higher level of resistance than the C6_00010W_A fusion, and we prioritized C4_02500C_A for further study (Table 1).

**Table 1.**
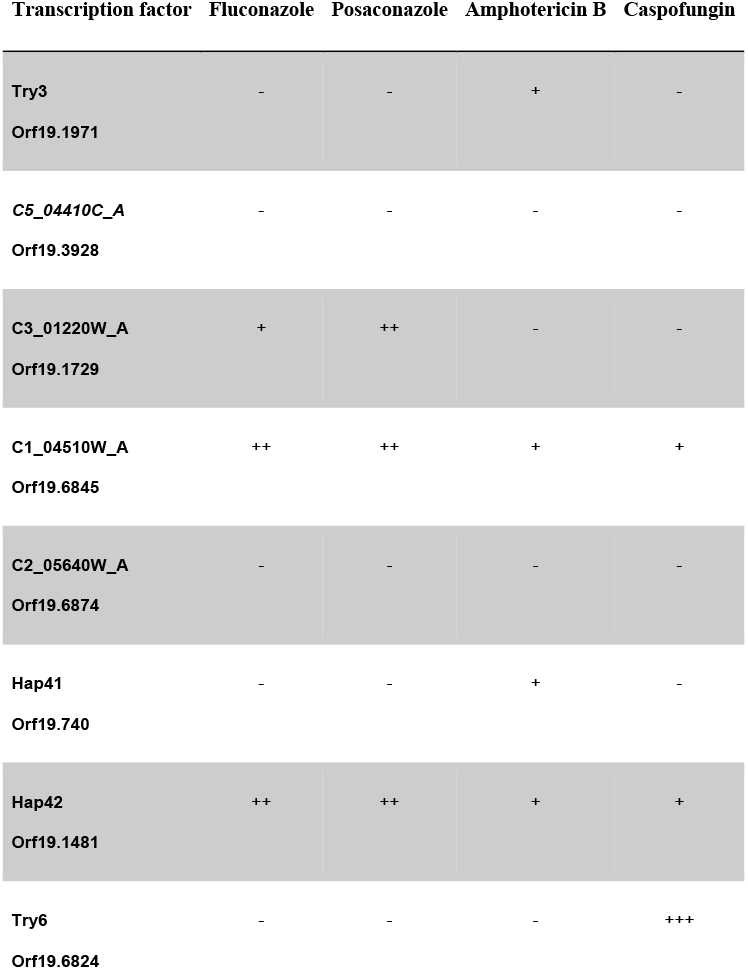
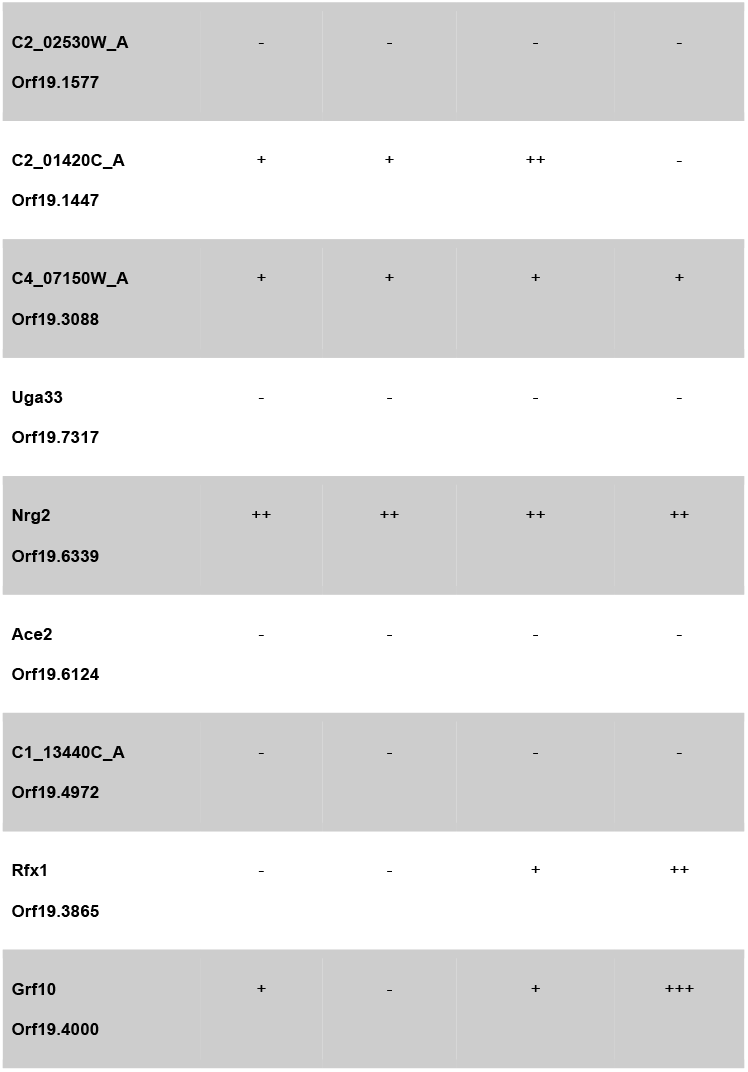
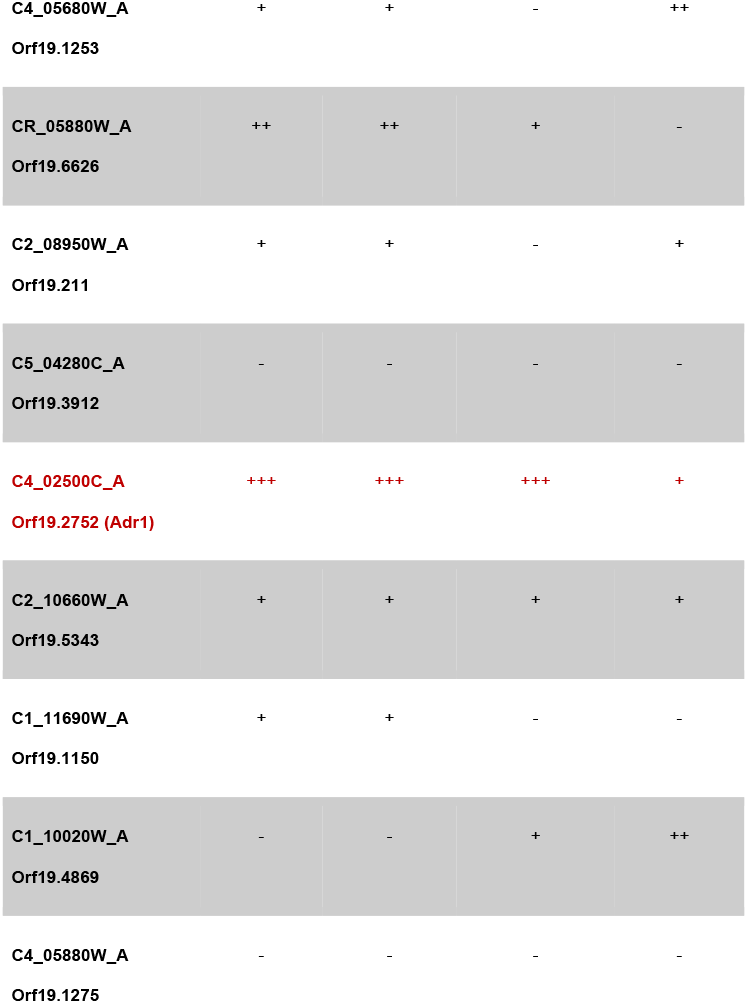
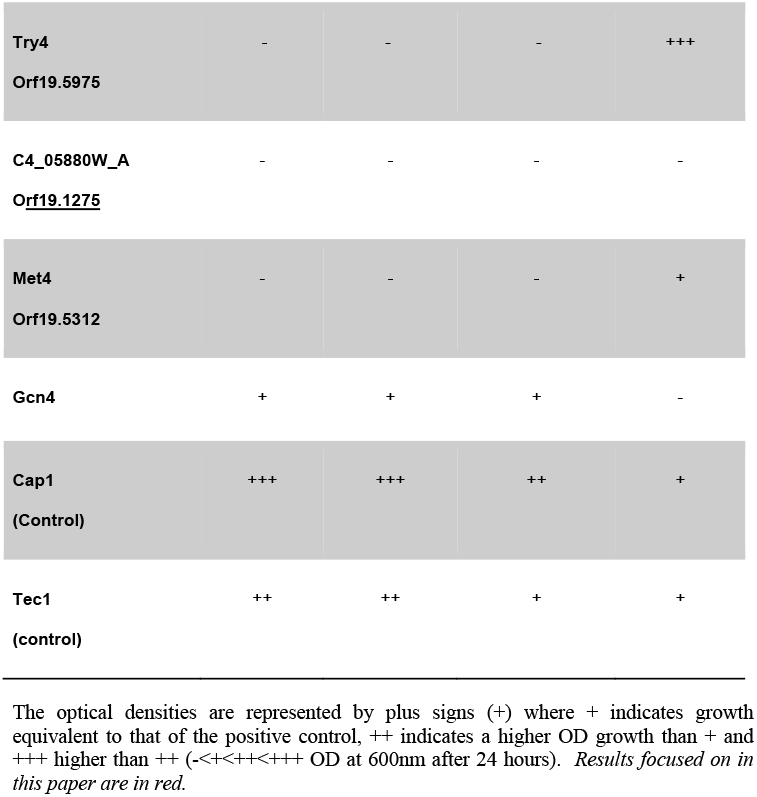
Screening of activated transcription factors in presences of different drugs

### Activation of C4_02500C_A confers multi-drug resistance

We tested whether the VP64 fusion to Orf19.2752 (C4_02500C_A) could trigger resistance to a variety of drugs - fluconazole, posaconazole, terbinafine, nystatin, caspofungin, anidulafungin and amphotericin B (Fig. 2A-B). The fusion of C4_02500C_A to VP64 increases the minimal inhibitory concentration as well as the minimal fungicidal concentration of fluconazole, amphotericin B and terbinafine (Fig. 2A), and also increased the MIC for these drugs (Fig. 2B), as well as to posaconazole and nystatin (not shown), by more than 3-fold. However, for the drugs caspofungin and anidulafungin that target the cell wall, there was no change in the MIC or growth rate for the activated strain relative to the control. Thus, activation of C4_02500C_A did not cause a general drug resistance but did seem effective in generating resistance to cell membrane targeting drugs.

**Fig. 2.**
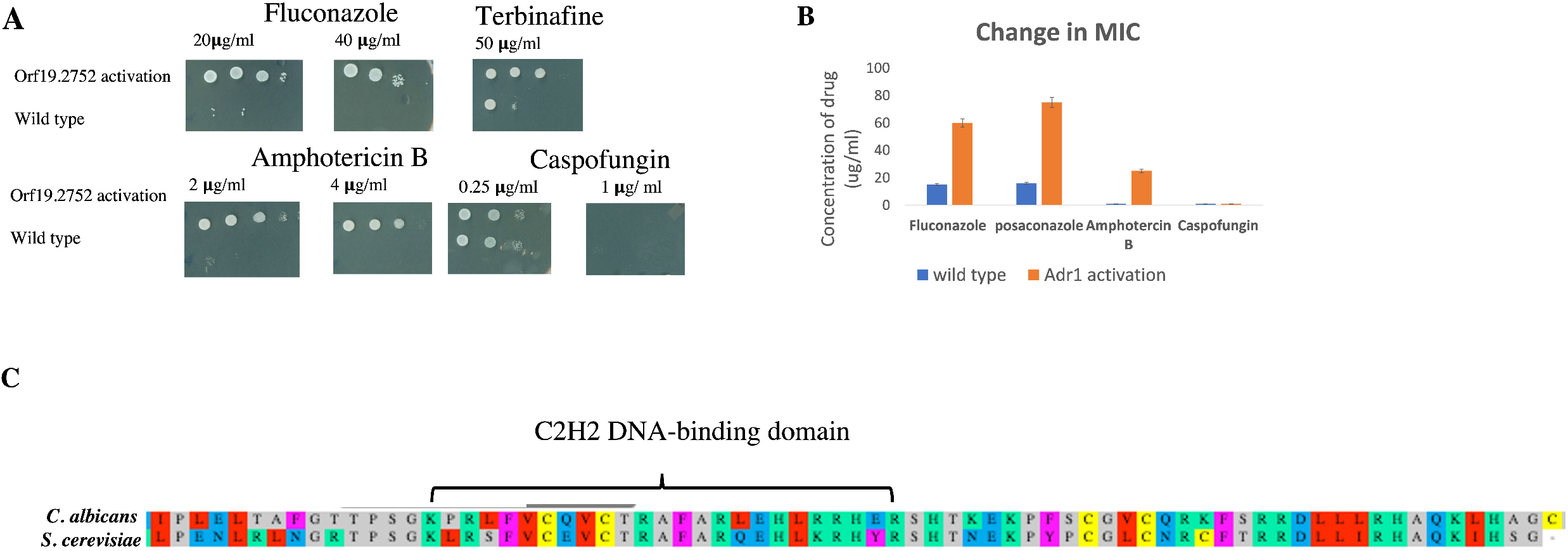
**(A) Plate assay of SC agar with different drugs confirming the VP64 fusion of Orf19.2752 generates fluconazole, terbinafine and amphotericin B resistance.** We used minimal inhibitory concentration (MIC) and 2MIC concentrations to check if the fusion construct creates resistance to the drugs. **(B) Activated Orf19.2752 caused fluconazole, posaconazole, terbinafine and amphotericin B resistance compared to wildtype.** Graphical representation of the change in minimal inhibitory concentrations of the Orf19.2752 activated strain in liquid SC media containing the various drugs. After 24 hrs cells were checked for residual growth activity on Sc media. **(C) Sequence alignment shows that Orf19.2752 is similar to the Adr1 protein of S. cerevisiae**. Blast alignment of the zinc cluster DNA binding domain of the Adr1 transcription factor in S. cerevisiae with that of Orf19.2752 in C. albicans.

### Orf19.2752 (C4_02500C_A) is an ortholog of *S. cerevisiae* Adr1

Because TFs are frequently conserved across species, we looked for orthologs of the *C. albicans C4_02500C_A* gene. We found it to be highly similar to the *S. cerevisiae ADR1* gene; the two proteins have about 50% sequence identity and the N-terminal DNA binding domain is highly conserved between them (Fig. 2C).

In *S. cerevisiae*, Adr1, acting through a conserved binding motif 5’ RCCCCM 3’, is required for transcriptional regulation of ethanol, glycerol and fatty acid utilization [25, 26]. Due to the highly conserved DNA binding domains of the two orthologs, we searched for this ScAdr1 binding motif upstream of *C. albicans* ORFs. We found 221 genes with this motif in their predicted promoter regions using Meme-suite software as described in the methods. Of the genes with this promoter motif, a significant fraction (one tenth, or 20 genes) was implicated in ergosterol biosynthesis (Fig 3A), while a further one quarter (52 genes) were categorized as of unknown function. However, in contrast to the situation in *S. cerevisiae*, this motif is not enriched in the ethanol, glycerol, and fatty acid metabolism genes in *C. albicans*. Because of the large number of motif-containing genes in the pathway of sterol biosynthesis, it appeared CaAdr1 might instead be linked to sterol production.

**Fig. 3.**
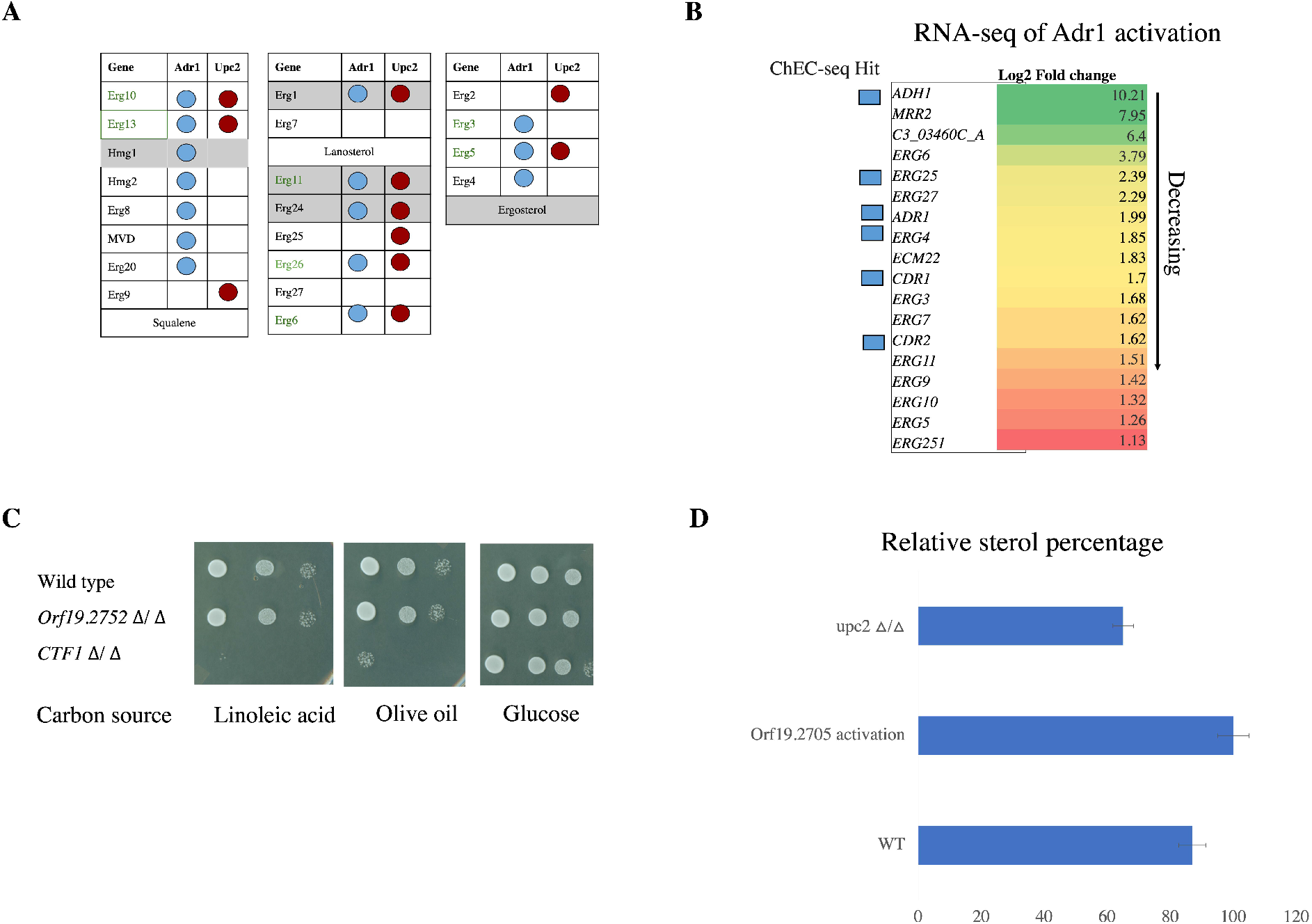
**(A) Presence of the Adr1 motif upstream of ergosterol genes.** The presence of the candidate Adr1 motif is represented by a blue circle. We found most of the ergosterol genes have the candidate Adr1 motif 5’NRCCCCM 3’ in their promoter regions. **(B) Transcriptomic profile of selected genes of the Orf19.2752-VP64 fusion strain shows upregulation of ergosterol genes.** After activation of the Adr1 transcription factor we did an RNAseq comparison of the activated strain and the wild type. We identified various ergosterol pathway genes upregulated and observed high expression of the genes for the transcription factor Mrr2 and the alcohol dehydrogenase Adh1. The full gene set is found in supplementary data file 2 along with the FPMK values. We also confirmed the binding of Adr1 by performing ChEC-Seq analysis; significantly bound genes are noted. **(C) Deletion of ADR1 (ORF19.2752).** We deleted the ADR1 gene and checked the resulting strain for fatty acid, glycerol and alcohol utilization. We found that it does not have any effect on growth on a fatty acid substrate, while a CTF1 deletion strain shows impaired growth on both linoleic acid or olive oil as a substrate. **(D) Sterol estimation of wildtype and the Orf19.2752 activated strain.** We extracted the sterols from overnight grown C. albicans by the pargolol-hexane extraction method. The extracted sterols showed a four-peak spectral absorption pattern characteristic of ergosterol and 24(28)-dehydroergosterol [24 (28)-DHE]. The activated strain showed an approximately 2-fold increase in measured sterols.

### Adr1 DNA binding motif

We further investigated potential 5’NRCCCCM3’ binding using ChEC-seq analysis; these results identified direct binding to several genes with the 5’NRCCCCM3’motif in their promoters, including Mrr2, Adh1, Ecm22, Erg5, Erg28, Cdr1, and Adr1. These genes were also upregulated in the Adr1-activation RNA-seq data set, suggesting that Adr1may be directly involved in transcriptional regulation of the ergosterol biosynthesis pathway (Fig. 3B). As well, the transcription factor Mrr2 is dramatically up-regulated; this could help explain the observed multi-drug resistance phenotype, as Mrr2 acts to up-regulate the *CDR* transporter-encoding genes, and our RNA-seq analysis shows both *CDR1* and *CDR2* are more highly expressed in the Adr1-VP64 fusion strain than in the control (Fig. 3B). To establish if the fluconazole resistance is a direct result of this upregulation of Mrr2, we deleted *MRR2* in the Adr1 activated strain. However, deletion of *MRR2* had essentially no effect on fluconazole resistance driven by activation of Adr1, suggesting the up-regulation of the Mrr2 TF was in fact not critical in creating the resistance phenotype (Fig. S1C).

### CaAdr1 influences sterol metabolism

Azole drugs target Erg11 of the ergosterol pathway in *C. albicans* [27], and upregulation of Erg11 is one of the known mechanisms for drug resistance against fluconazole [28, 29]. To test if the *ADR1* gene of *C. albicans* was involved in sterol metabolism, we deleted the gene and checked the consequences of loss of function; consistent with a role in sterol biosynthesis, *ADR1* deletion causes slight sensitivity to cell membrane targeting drugs like fluconazole (Fig. 2A; Fig. S1 D). The complementation of *adr1* with the native protein or the Vp64 activated version restored the drug sensitivity (Fig. SD). However, unlike the situation with *ScADR1*, the deletion of *CaADR1* did not block growth on fatty acid substrates (Fig. 3). By contrast, deletion of the gene for the transcription factor Ctf1, identified as a regulator of fatty acid metabolism genes in the pathogen [30], completely blocked *C. albicans* growth on linoleic acid and severely compromised growth on olive oil (Fig. 3C). This suggests that in *C. albicans* Ctf1 is controlling fatty acid utilization, while Adr1 is not involved. In *S. cerevisiae*, ScAdr1 was found to be haplo-insufficient for ethanol, glycerol and fatty acid metabolism.

Similarly, in *C. albicans*, CaAdr1 is haplo-insufficient for fluconazole resistance, as the heterozygote showed sensitivity to fluconazole relative to the WT but was clearly more resistant than the homozygous null (Fig. S1C).

We directly checked the sterol content of the Adr1 activated strain and wild-type by extracting sterols with the organic solvents pargolol and hexane followed by spectrophotometric assessment. Activated CaAdr1 enhances the production of ergosterol (Fig. 3D). We also directly assessed the transcriptional consequences of Adr1 activation through RNAseq analysis. *ADH1*, which encodes the alcohol dehydrogenase that oxidizes ethanol to acetaldehyde [31] was the most highly upregulated gene in our profile, and intriguingly, the orthologous gene in *S. cerevisiae* is a direct target of ScAdr1.

### Adr1 and Upc2 roles in azole response

Consistent with the presence of the candidate Adr1 binding motif in their promoters, we found the expression level of most of the ergosterol pathway genes, including Erg11, was upregulated by the activated Adr1 construct. This increase in ergosterol pathway gene expression was however not associated with up-regulation of the classic ergosterol biosynthesis pathway transcription factor Upc2, which functions as a key regulator of the pathway in both *S. cerevisiae* and *C. albicans* [32–34]. Therefore, it appears that Adr1 activation of the *C. albicans* ergosterol pathway genes is likely direct (Supplementary Table 1), and thus the fluconazole resistance generated by the Adr1VP64 fusion protein may be due to the generalized increase in the expression of these ergosterol biosynthetic pathway genes. We then asked whether the fluconazole resistance generated by Adr1 activation was fully independent of Upc2. First, we compared fluconazole resistance levels in strains with Adr1 activated and with Upc2 activated, as well as in strains with deleted *UPC2* or *ADR1*. As is shown in Fig. 4A-B, both Upc2 activation and Adr1 activation gave similarly high levels of resistance to fluconazole, while both Adr1 deletion and Upc2 deletion conferred sensitivity to fluconazole, with the Upc2 deletion strain being somewhat more sensitive. Secondly, we assessed the resistance to fluconazole of the doubly activated strain; in this case the strain grew poorly in the absence of drug but was resistant to fluconazole at similar levels to that of the single activated mutants (Fig. 4A-B). Finally, we investigated the behaviour of cells with activated Adr1 that lacked functional Upc2, and cells with an activated Upc2 that lacked Adr1. We observed that loss of Upc2 had essentially no effect on the fluconazole resistance caused by activation of Adr1, suggesting the effect of Adr1 on drug resistance is independent of Upc2, while loss of Adr1 significantly compromised fluconazole resistance caused by activation of Upc2 (Fig. 4A-B). This is consistent with part of the effect of Upc2 activation on azole resistance working through Adr1. We checked the upstream sequences of the *UPC2* and *ADR1* genes for regulatory motifs and found that the promoter sequence of the *ADR1* gene contains a potential *UPC2* motif (Fig. 4C), consistent with Adr1 being part of the Upc2 regulon, while the *UPC2* promoter lacks any potential Adr1 binding motif.

**Fig. 4.**
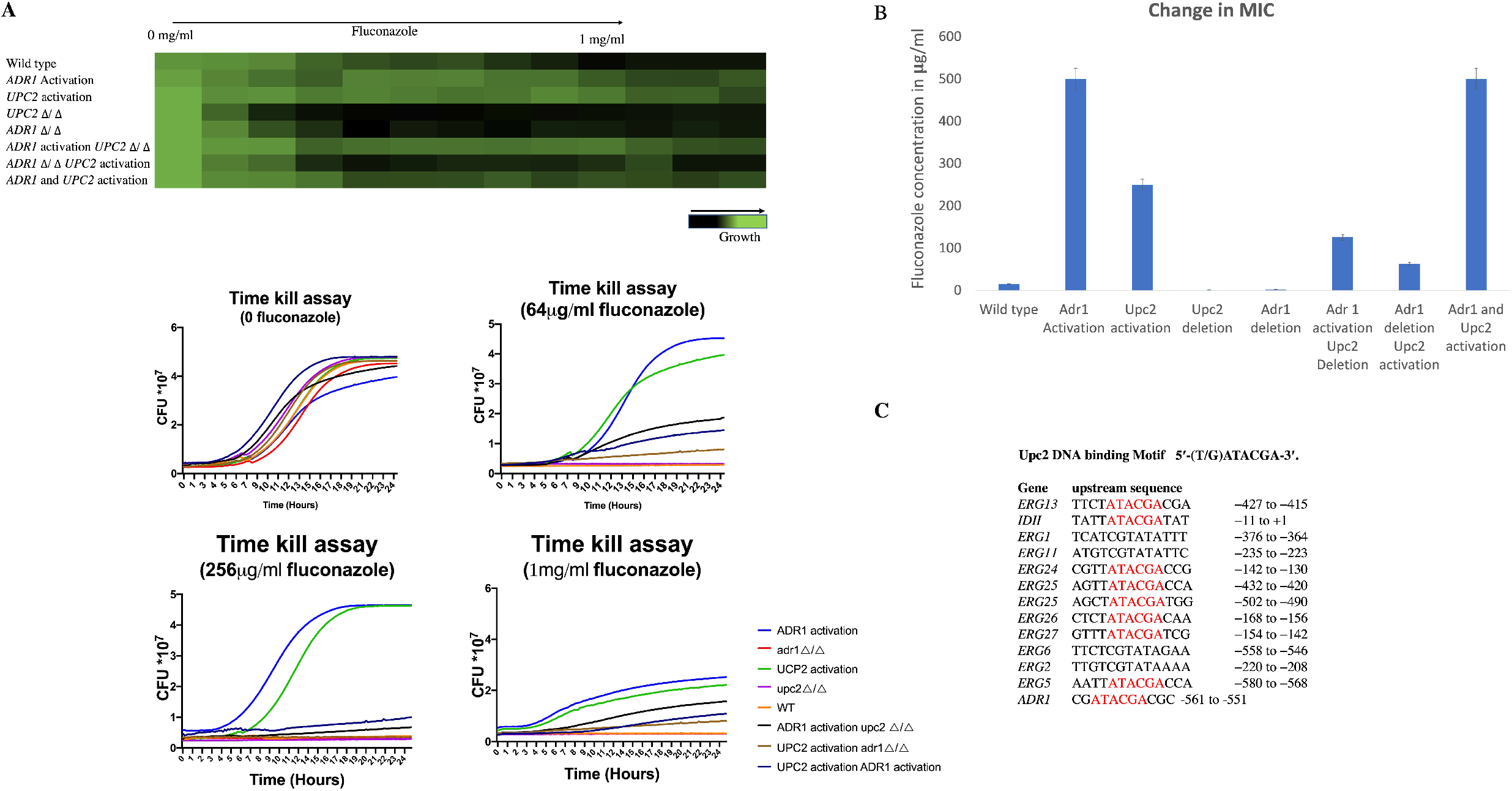
**(A) Adr1 and Upc2 influence response to fluconazole**. Fluconazole MIC assay of ADR1 deletion, ADR1 activation ADR1 activation in UPC2 deletion, ADR1 deletion in UPC2 activation and double activation, followed by the time kill assay. **(B) Fold change in MIC from Adr1 and Upc2 activation and deletion**. Graphical representation of the change in minimal inhibitory concentrations of various drugs in the Orf19.2752 activated strain. **(C) Upc2 DNA binding motif upstream of various regulons including ADR1.** Upstream regions of genes including the transcription factors Adr1, Upc2, Mrr2 and Mrr1 as well as the structural genes for the ergosterol biosynthesis pathway were examined for candidate TF binding motifs. We identified a potential Upc2 binding motif upstream of the ADR1 gene.

### Phylogenetic analysis

We examined a phylogenetic profile of the Ascomycota and characterized the upstream sequences of all ergosterol biosynthesis genes for the regulatory motifs associated with both the Adr1 and the Upc2 transcription factors. In *C. albicans* our data suggest that Adr1 works together with Upc2 to control the ergosterol biosynthesis pathway, with Adr1 also controlling the expression of genes such *ADH1*, encoding the alcohol dehydrogenase which oxidizes ethanol to acetaldehyde, and *MRR2*, encoding a stress-responsive transcription factor. The more basal filamentous fungi also have the candidate binding motifs for both Adr1 and Upc2 the promoters of the ergosterol biosynthesis genes. However, the presence of the Adr1 motif in the promoter of *MRR2* is very specific to the CTG clade species (Supplementary data file 2).

In the evolutionary trajectory leading to *S. cerevisiae*, it appears that the Adr1 TF was repurposed to control alcohol, fatty acid and glycerol utilization, taking over from Ctf1 orthologs that do the job in the filamentous fungi and the CTG clade species. Based on the search of promoter motifs, we uncovered that Upc2 motif is found in the promoters of ergosterol pathway genes throughout the fungi along with the Adr1 motif. But after *C. guilliermondii* in the phylogeny, the Adr1 motif signal weakens, leaving only Upc2 with a strong signal associated with the ergosterol pathway. The Adr1-binding motif signal connected the ergosterol pathway genes in the CTG clade appears to be gradually transferred to genes involved in control of alcohol and fatty acid utilization in *S. cerevisiae* and its relatives (Supplementary data sheet 1).

## Discussion

One of the common approaches to investigating the regulatory networks controlling drug resistance in fungal pathogens is through comparison with the *S. cerevisiae* circuits; this approach has led to the discovery of many TFs responsible for drug resistance in both *S. cerevisiae* and *C. albicans* including Fcr1, Ndt80, Mrr1 and Upc2 [32, 35–38]. However, these two fungi diverged as long ago as 300 million years, and for species diverged by such an evolutionary distance, the majority of the DNA-binding patterns of a given regulator in one species are unlikely preserved in the other species [20]. Overall, genome-wide correlations converge on about 10% overlap for species with this level of divergence [20], and therefore, there is a significant probability that many of the TFs responsible for drug resistance will be different between *C. albicans* and the budding yeast. To identify candidate TFs with Candida-specific roles in drug resistance, we selected a set of TF-encoding genes that were either not found in *S. cerevisiae* or that were predicted to have potentially changed function between the two species. We identified 30 such TFs and activated them in order to identify potential roles in drug resistance (as well as other cellular processes) (Table 1). Among these transcription factors, Orf19.2752 activation resulted in clear resistance to a set of drugs generally targeting the cell membrane; activation of this transcription factor generated resistance to azoles, allylamines and polyenes. Sequence comparisons established that *C4_02500C_A (ORF19.2752)* was the *C. albicans* ortholog of the *S. cerevisiae* gene *ADR1*, a gene not linked to drug resistance in the budding yeast. These two transcription factors share a highly conserved DNA binding domain (Fig. 1C).

In *S. cerevisiae*, Adr1 is involved in the transcriptional regulation of genes involved in the catabolism of ethanol, glycerol and fatty acids [25, 26], and is proposed to act through a candidate DNA-binding motif, 5’NRCCCCM3’ in the promoter regions of these genes [39]. Interestingly, in *C. albicans* this same DNA-binding motif is enriched in the upstream regions of the ergosterol-pathway genes, whereas it was generally absent from the promoters of the ethanol, glycerol and fatty acid utilization genes of this pathogen. This suggests that Adr1 may have been rewired from the ergosterol biosynthesis pathway in other fungi to the metabolic utilization of ethanol, glycerol and fatty acids in *S. cerevisiae*. Further investigation established that activation of CaAdr1 generated transcriptional upregulation of most of the ergosterol pathway genes in the pathogen. However, Upc2, the well-established regulator of the Erg-pathway genes in both *C. albicans* and *S. cerevisiae*, was not transcriptionally upregulated, suggesting that in *C. albicans* Adr1 activation of the ergosterol pathway genes was not going through Upc2.

To further investigate the proposed Adr1 DNA binding motif, we performed a ChEC-seq analysis [40] and found several the drug-resistance-related genes including Mrr2, Adh1, Ecm22, Erg5, Ddr48, Erg28, Cdr1 and Adr1 were both up-regulated in the RNAseq analysis and were ChEC-seq hits. Previous *in-silico* analysis of a number of TFs had shown a weak but statistically significant overlap in the genes in *S. cerevisiae* and *C. albicans* containing the Adr1 motif in their promoters [41]. This is consistent with our ChEC-seq analysis; while many genes have rewired between the two species, some genes, like *ADH1*, are under Adr1 control in both species, suggesting that Adr1 might have multiple roles in both fungi. However, the bulk of the circuit of ethanol, fatty acid and glycerol metabolism controlled by Adr1 in *S. cerevisiae* is under the control of Ctf1 in *C. albicans* [30], as Adr1 deletion did not affect growth on substrates like olive oil and linoleic acid, whereas Ctf1 deletion gives a clear auxotrophy.

These data underline the multiple circuit restructurings the control of these pathways in different fungi. In *S. cerevisiae* Cat8 and Adr1 both appear to have rewired to connect to the module controlling alcohol, acetic acid and fatty acid utilization; Adr1 from the ergosterol circuit, Cat8 from gluconeogenesis (ScSip4). Another event is the disappearance of the Ctf1 TF from the *S. cerevisiae* genome, as there is no apparent ortholog of Ctf1 in *S. cerevisiae*. This loss could be facilitated by transfer of the Ctf1 regulon to Adr1 control in *S. cerevisiae*. In *S. cerevisiae* Upc2 gains a paralog (Emc22) and apparently unique control over the ergosterol pathway [42].

In *C. albicans*, Adr1 activation causes upregulation of many ergosterol pathway members, Erg11 (target of azoles), Erg1 (target of allylamines), Erg2 (target of morpholines), HMG1/2 (target of statins), and causes increases in ergosterol itself (target of polyenes), which has the potential of generating multidrug resistance. The activation of Adr1 dramatically enhances the MIC of fluconazole, amphotericin B, terbinafine and statins. We created a series of strains in order to determine the how Upc2 and Adr1 are influencing the ergosterol pathway. Activation of either Upc2 or Adr1 enhanced azole resistance, while deletion of either gene created azole sensitivity. Activation of both TFs at the same time caused poor growth, perhaps due to the disturbed circuits or due to the activation of an Erg3-driven side branch of the pathway generating the toxic 14a methyllergosta 8-24(28) dienol. However, the relative resistance to azoles remained similar to the individually activated strains, suggesting the actions of the two TFs are not additive or synergistic. Another known stress associated with ergosterol is hypoxia, and therefore we tested both Adr1 activation and *adr1* deletion under hypoxia conditions [43, 44]. We did not find any changes in the *adr1* deletion strain under hypoxia, while the up-regulated allele somewhat improved growth (Fig. S1 E). Previously, the *upc2* deletion strain has been reported to show a significant growth defect under hypoxia [45], so this distinction will need further investigation.

The fluconazole resistance caused by activation of Upc2 is significantly dependent on the presence of Adr1, but loss of Upc2 had very little effect on the resistance profile of strains with an activated Adr1. In addition, there is a potential Upc2 DNA binding site in the promoter region of *ADR1*, while *UPC2* has no candidate site for Adr1. These results are consistent with Upc2 serving as a master regulator of ergosterol biosynthesis, controlling ERG gene expression both directly and through regulation of the Adr1 TF that also serves as an activator of ERG gene expression. In the absence of Upc2, Adr1 is sufficient to ensure ERG expression, although response to azole drugs is compromised by loss of either TF. A strain containing deletions of both *adr1* and *upc2* was very slow growing, and also very sensitive to fluconazole.

Among the highly up-regulated genes resulting from Adr1 activation is the gene encoding Mrr2, itself a TF involved in the expression of the multi-drug-resistance regulating membrane transporter Cdr1. However, even though *CDR1* expression was somewhat up-regulated in the Adr1 activated strain, the Mrr2 up-regulation did not seem critical for the observed fluconazole resistance, because deletion of *MRR2* in the Adr1 activated strain did not compromise this resistance.

While it appears that in filamentous fungi and the CTG clade species Adr1 is linked to ergosterol biosynthesis, in the *Saccharomycotina* it has been shifted to control the pathway for ethanol, glycerol and fatty acid utilization [46, 47] replacing Ctf1 that controls the process in the *non-Saccharomycotina* species. This transfer appears so complete that the Ctf1 factor has been lost in the *Saccharomycotina*. Intriguingly, throughout this transition, Adr1 has maintained a role in the control of the expression of the alcohol dehydrogenase catalyzing the oxidation of ethanol to acetaldehyde (*ADH1* in *C. albicans, ADH2* in *S. cerevisiae*). To deal with ethanol toxicity, in *S. cerevisiae* ethanol is modified into unsaturated lipids and ergosterol [48]. The rewiring of Adr1 from the ergosterol pathway to the ethanol utilization process [39, 49] may have been driven by the shift to the Crabtree positive lifestyle of *S. cerevisiae*, requiring the ability to both tolerate and process high concentrations of ethanol.

## Conclusion

Sterol biosynthesis is critical for fungal biogenesis and a target for many antifungal drugs. We have identified the TF Adr1 as a key regulator of sterol biosynthesis in the fungal pathogen *C. albicans*, where it works in concert with the zinc cluster TF Upc2. We suggest Upc2, when bound to ergosterol, remains inactive in the cytoplasm, but is activated when there is depletion of ergosterol, whether by environmental factors or due to the presence of the drugs targeting the ergosterol pathway. Activated Upc2 goes to the nucleus and turns on key players of the ergosterol biosynthetic pathway as well as the TFs Mrr1 and Adr1. Adr1 aids in regulating the ergosterol pathway genes and turns on the TF Mrr2. Thus, activation of Upc2 and Adr1 leads to the co-ordinated expression of the ergosterol biosynthesis pathway, as well as the activation of phospholipid transporters Cdr1 and Cdr2 that can also function to export antifungal drugs (Fig. 5).

**Fig. 5.**
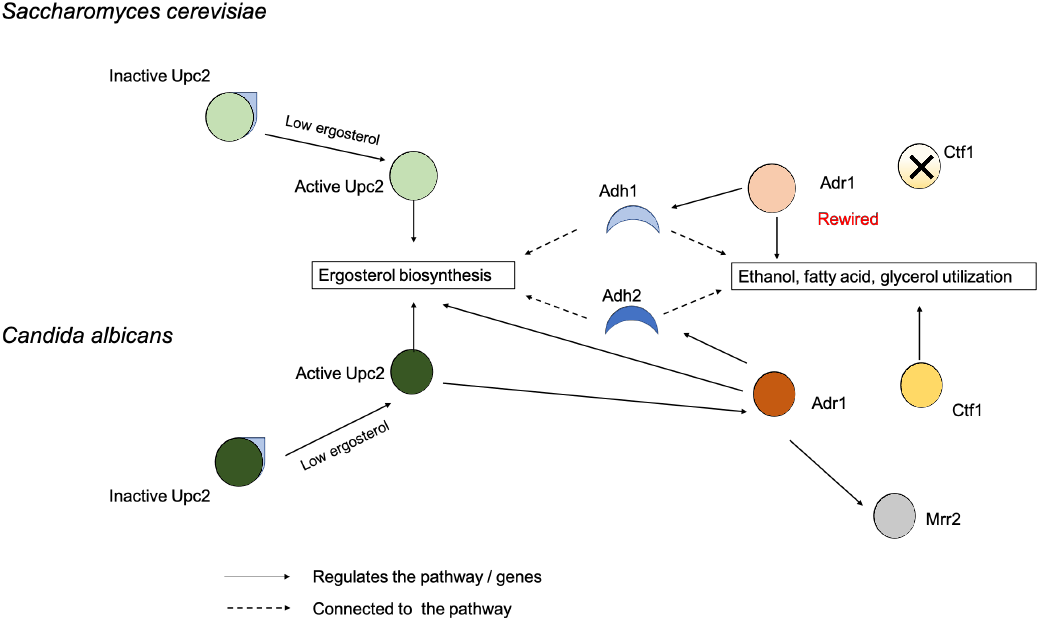
Proposed model of the multiple rewiring of transcription factors between S. cerevisiae and C. albicans along with coordination of Upc2, Adr1 and Mrr2 during the presence of azoles or depletion of ergosterol leading to drug resistance in C. albicans. In both S. cerevisiae and C. albicans, the Upc2 transcription factor binds to ergosterol and remains inactive in cytosol. Depletion of ergosterol changes Upc2 to an active form, which enters the nucleus and initiates the transcription of the genes for ergosterol biosynthesis. In C. albicans activated Upc2 also triggers expression of the Adr1 transcription factor which further serves to direct expression of ergosterol biosynthesis genes, as well as the alcohol metabolism gene Adh1 and the Mrr2 transcription factor. In S. cerevisiae, Adr1 has been rewired to control other parts of the alcohol utilization circuit in addition to alcohol dehydrogenase, as well as to control both fatty acid and glycerol utilization circuits.

## Materials and Methods

### Bioinformatics analysis

Sequences of genes *CaADR1and ScADR1* were obtained from the Candida Genome Database (CGD-http://www.candidagenome.org/) and the Saccharomyces Genome Database (https://www.yeastgenome.org/). Gene orthogroup assignments for all predicted protein-coding genes across 23 ascomycete fungal genomes were obtained from the Fungal Orthogroups Repository [50] maintained by the Broad Institute (broadinstitute.org/regev/orthogroups).

DNA sequence motifs were identified using the Web-based motif-detection algorithm MEME (http://meme.sdsc.edu/meme/intro.html; [51]. For more stringent motif identification, we used MAST hits with an E value of <50. An E value of 500 corresponds roughly to a p-value of 0.08 in our analysis, and an E value of 50 roughly corresponds to a p-value of 0.008. We also used AME (http://meme-suite.org/tools/meme), which identifies known motifs throughout the Candida upstream sequences.

Protein domains and linear motifs were detected from each individual TF protein sequence using INTERPROSCAN, PFAM and ELM motif definitions. For ChEC-seq analysis we used Integrative Genomics Viewer (IGV) (https://igv.org) and MEME-ChIP software.

### Strains and culture conditions

For general growth and maintenance of the strains, the cells were cultured in fresh YPD medium (1 % w/v yeast extract, 2 % w/v Bacto peptone, 2 % w/v dextrose, 80 mg/L uridine with the addition of 2 % w/v agar for solid medium) at 30 °C. For drug essays we used synthetic dextrose (SD) medium (0.67 % w/v yeast nitrogen base, 0.15 % w/v amino acid mix, 0.1 % w/v uridine, 2 % w/v dextrose and 2 % w/v agar for solid media) along with the various concentrations of drugs in liquid and solid media.

### Gene knockouts using CRISPR

All *C. albicans* mutants were constructed in the wild type strain CAI4. The protocol used for the CRISPR-mediated knockout of *ADR1, CTF1* and *UPC2* was adapted from [52]; we used *URA3* replacements in our study. CRISPR-mediated knockouts used the lithium acetate method of transformation with the modification of growing transformants overnight in liquid YPD at room temperature after removing the lithium acetate-PEG. *C. albicans* transformants were selected on SD-URA plates.

### Activation of transcription factors

For the activation module, the *ACT1* promoter and VP64 were amplified by PCR and homology was created by primer extension such that there is an Mlu I site between *ACT1* and VP64. After this, the C-terminus of the VP64 cassette was cloned into the multiple cloning site. pCIPACT and the ligated Act1-VP64-MCS were digested with Hind III and were ligated using T4 DNA ligase. This ligated CIPACT-VP64 plasmid was transformed into *E. coli* using the calcium chloride method. High-throughput equipment at the Concordia Genome Foundry, the Biomek FXP Automated Workstation otherwise known as Biomek FXP liquid handler was used to create and screen this library.

Plasmids extracted from colonies that were determined to have the guide sequence successfully cloned in were then used to transform *C. albicans* using a lithium acetate transformation protocol. pCIPACT1 was linearized by StuI-HF digest, and 1-2 μg of the linearized plasmid was used in the transformation. *C. albicans* transformants were selected on SD URA-plates.

### RNA seq analysis

The CaAdr1 and SC5143 strains were grown in SC media overnight at 30 °C, diluted to OD600 of 0.1 in YPD at 30 °C, and then grown to an OD_600_ of approximately 1.0 on a 220-rpm shaker. Total RNA was extracted from 2 biological replicates using the Qiagen RNeasy Minikit protocol, and RNA quality and quantity were determined using an Agilent bioanalyzer. Paired-end sequencing (150bp) of extracted RNA samples was carried out at the Quebec Genome Innovation Center located at McGill University using an Illumina miSEQ sequencing platform. Raw reads were pre-processed with the sequence-grooming tool cutadapt version 0.4.1 [53] with the following quality trimming and filtering parameters (‘--phred33 --length 36 -q 5 --stringency 1 -e 0.1’). Each set of paired-end reads was mapped against the *C. albicans* SC5314 haplotype A, version A22 downloaded from the Candida Genome Database (CGD) (http://www.candidagenome.org/) using HISAT2 version 2.0.4. SAM tools was then used to sort and convert SAM files. The read alignments and *C. albicans* SC5314 genome annotation were provided as input into StringTie v1.3.3 [54], which returned gene abundances for each sample. Raw and processed data have been deposited in NCBI’s Gene Expression Omnibus [55].

### Sterol quantification

The cells were grown overnight (16 hrs) at 30°C and harvested by centrifugation. Non-treated cells were maintained separately and considered as control. The cell pellets were washed with sterile distilled water twice. We followed the same method which has been described in [56] with slight modifications. Cell pellets were resuspended in 2.5 ml methanol, 1.5 ml potassium hydroxide (60% [wt/vol]), and 1 ml methanol-dissolved pyrogallol (0.5% [wt/vol]) and heated at 90°C for 2 h. The cell extracts were cooled and then sterols were extracted with two rounds of treatment with 5ml of hexane. The extracted sterols indicated a four-peak spectral absorption pattern produced by ergosterol and 24(28)-dehydroergosterol [24 (28)-DHE] spectrophotometrically (DU530 life science UV spectrophotometer). Both ergosterol and 24(28)-DHE absorb at 281.5 nm, whereas only 24(28)-DHE absorbs at 230 nm. The ergosterol content is determined by subtracting the amount of 24(28)-DHE (calculated from the A230) from the total ergosterol plus-24(28)-DHE content (calculated from the A281.5). Ergosterol content was calculated as a percentage of the wet weight of the cells with the following equations: % ergosterol + % 24(28)-DHE [(A281.5/290) * F]/pellet weight, % 24(28)-DHE _ [(A230/518) _ F]/pellet weight, and % ergosterol = [% ergosterol + % 24(28)-DHE] – [% 24(28)-DHE], where F is the factor for dilution in petroleum ether and 290 and 518 are the E values (in percent per centimeter) determined for crystalline ergosterol and 24(28)-DHE, respectively.

### ChEC-seq analysis

To perform the ChEC-seq analysis we followed [40]. We constructed Adr1-MNase strain and grow the overnight cultures of *C. albicans* Adr1-MNase tagged and free MNase strains, then they were diluted to a starting OD600 of 0.1 in 50 ml YPD medium and grown at 30°C to an OD600 of 0.7 to 0.8. Cells were washed three times with 1 ml buffer A (15 mM Tris [pH 7.5], 80 mM KCl, 0.1 mM EGTA, 0.2 mM spermine, 0.5 mM spermidine, one tablet Roche complete EDTA-free mini protease inhibitors, 1 mM phenylmethylsulfonyl fluoride [PMSF]) and were then resuspended in 800 μl buffer A containing 0.1% digitonin (Sigma) and permeabilized for 10 min at 30°C with shaking. MNase digestions were performed by adding CaCl2 to a final concentration of 5 mM and incubated for different time points at 30°C. At each time point, a total of 200-μl aliquots of the ChEC digestions were transferred to a tube containing 50 μl of 250 mM EGTA to quench MNase digestion. DNA were extracted using MasterPure yeast DNA purification kit (Epicentre, MPY80200). ChEC DNA was subjected to size selection using the Pippin Prep (SageScience) size selection system with a 2% agarose gel cassette, allowing the removal of multikilobase genomic DNA fragments and the enrichment of 100- to 400-bp DNA fragments.

## Acknowledgements

We thank all members of M.W lab, Dr. Smita Amarnath and Dr. Nicholas Gold. for valuable comments during this study. We want to acknowledge funding support from the NSERC Discovery RGPIN/4799 (M.W), and NSERC Canada Research Chair 950-228957 (M.W) awards. We would also like to thank Concordia University NSERC Synbioapp program and bio-foundry. A.S. is supported by a Fonds de Recherche du Québec - Santé J2 salary award. We would like to thank Faiza Tebjji, CT Anagha and Maria Mantilla for guiding and helping me for the CheC-seq experiment and sharing their lab facilities. We also want to thank Dr. Joachim Morschhäuser for sending us *C. albicans* strains for this project.

## Figure legends

**Supplementary figure 1.**
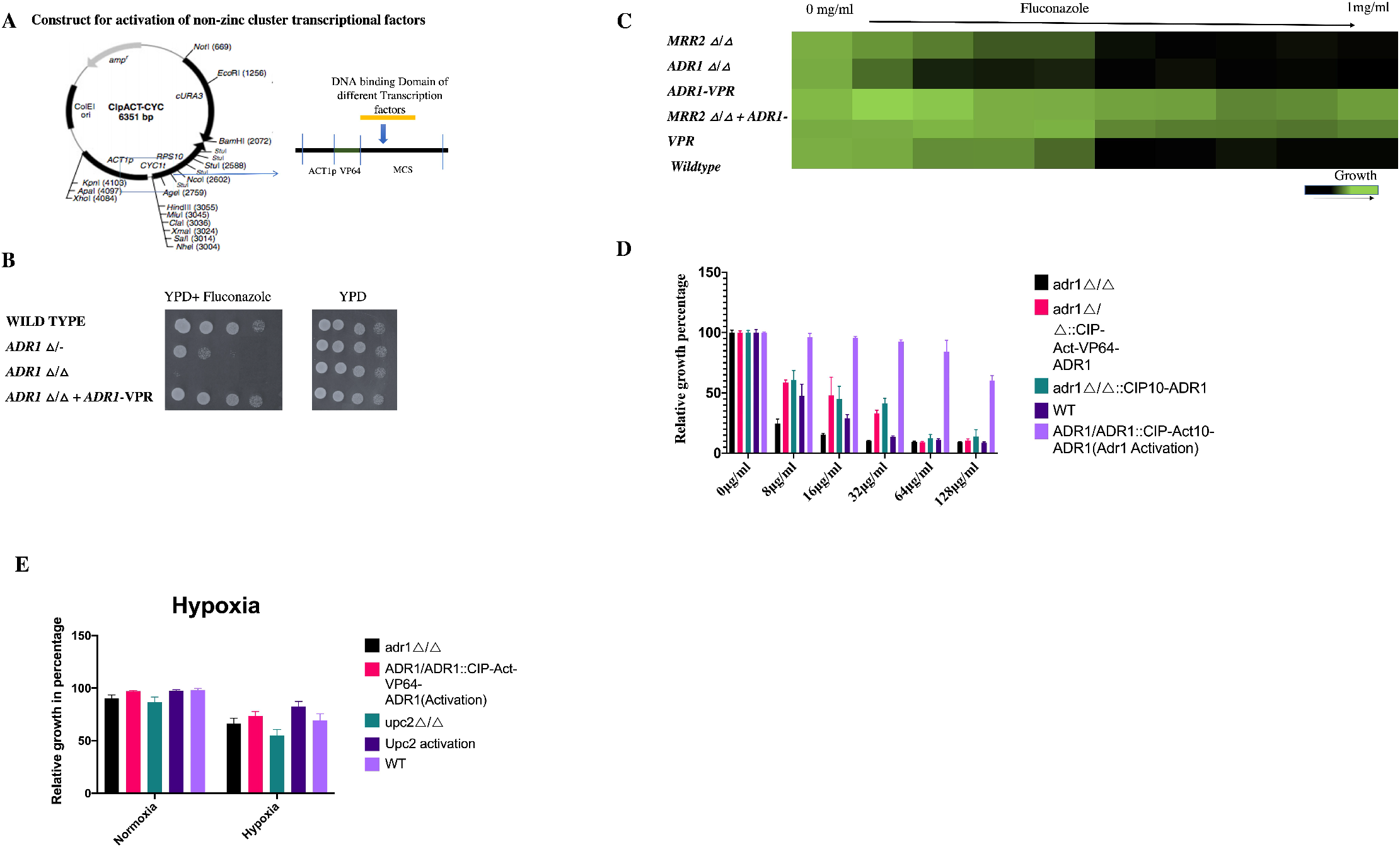
(A) The construct for activation of non-zinc cluster transcription factors. (B) The adr1Δ/Δ strain (ORF19.2752) showed fluconazole sensitivity. The heterozygous adr1Δ/+ shows better growth than the adr1Δ/Δ in the presence of fluconazole. (C) The mrr2 Δ/Δ in the Adr1 activated strain caused no changes in drug resistance. (D) The graphical representation of the growth variation in presence of different concentrations of fluconazole in the adr1Δ/Δ, adr1Δ/Δ with complementation of hyperactivated ADR1(Adr1-VP64) allele, adr1Δ/Δ with complementation of the native ADR1 allele, Adr1 hyperactivation (Adr1-VP64) and wildtype. (E) The graphical representation of the growth variation under Hypoxia in the adr1Δ/Δ, Adr1 hyperactivation (Adr1-VP64), Upc2Δ/Δ, Upc2 hyperactivation and wildtype.

